# Factors determining success of the chronically instrumented unanesthetized fetal sheep model of human development: a retrospective cohort study

**DOI:** 10.1101/2022.06.17.496637

**Authors:** Colin Wakefield, Mingju Cao, Patrick Burns, Gilles Fecteau, Andre Desrochers, Martin G Frasch

**Affiliations:** Drexel University College of Medicine, Philadelphia, PA, USA; CHU Ste-Justine Research Center, Montreal, QC, Canada; Dept. of Clinical Sciences, School of Veterinary Medicine, University of Montreal, St-Hyacinthe, QC, Canada; Dept. of Obstetrics and Gynecology and Dept. of Neurosciences, University of Montreal, Montreal, QC, Canada; CRRA, School of Veterinary Medicine, University of Montreal, St-Hyacinthe, QC, Canada; Dept. of OBGYN, CHDD, University of Washington, Seattle, WA, USA

**Keywords:** fetal sheep model, chronic instrumentation, outcomes, retrospective study

## Abstract

**Aim:** Chronically instrumented non-anesthetized fetal sheep (CINAFS) have been the mainstay model of human fetal development for 60 years. As a large “two for one” animal model, with instrumentation of the ewe and her fetus, the model poses challenges to implement de novo and to maintain overtime at the highest standards of operating procedures to ensure its ongoing performance. A common, yet conventionally underreported issue researchers face is the rate of animal loss. Here, we investigate what determines the success of the pregnant sheep model.

**Methods:** We conducted a retrospective cohort study consisting of 82 experiments spanning the course of six years. Our team identified ten variables that we anticipated were likely to influence the experimental outcome, such as the time of year, animal size, and surgical complexity.

**Results:** The single variable identified in this study as determining the successful outcome of the experiments is the experience level of the team.

**Conclusion:** The CINAFS model offers enormous potential to further our understanding of human fetal development and to create interventional technologies. However, to improve the outcomes of CINAFS models, improved communication and training are needed. We discuss the implications of our findings for the successful implementation of this challenging yet scientifically advantageous animal model of human physiology.

**Key points:**

- The fetal sheep model closely mirrors the physiology of human fetal development
- In addition to its high translational potential, this model is known to have some generally not reported rate of experimental failure
- We show that factors such as animal characteristics & surgical complexity do not influence the experimental outcomes
- Instead, the key factor in model experimental success is the level of the research team’s experience
- The key factors to improve the animal model outcomes are an intra- and inter-team communication

## Introduction

Of the animal models of fetal physiology currently in use, fetal sheep model stands out as having been demonstrated over 60 years to best mimic human physiology.^1–3^

Prenatal medicine has adopted the use of a pregnant sheep model for obvious reasons of the preclinical modeling of health and disease as well as to overcome the fact that researchers cannot directly monitor the human fetus. Limitations in access to the human fetus and considerable risk associated with sampling, make human testing nearly impossible. Noninvasive (e.g., cardiotocography) and minimally invasive (e.g., chorionic villus sampling) techniques to assess human fetal physiology and development exist, however, they are largely limited to informing clinical care in high-risk human pregnancies. These technologies offer minimal insight into the physiological, endocrine, and metabolic development of the fetus.^1^ This lack of direct access to monitoring the fetal condition has limited our understanding of fetal development and the impact of biological & environmental factors on postnatal health outcomes.

The resulting gap in knowledge of fetal development is an enormous issue for the medical community. Worldwide, 12% of childbirths are premature, two-thirds of which occur without a known risk factor. Premature childbirth accounts for the majority of deaths globally in children under five.^4^ In addition, children who survive premature birth are at increased risks for respiratory, immune, neurologic, feeding, hearing, neurodevelopment, behavioral, socio-emotional, and learning complications. Seemingly benign factors during fetal development, such as maternal stress, significantly influence the life-long health trajectories of the fetus.^5,6^

The pregnant sheep model is an ideal candidate for recreating human development in an experimental setting, allowing researchers to investigate human development. There are several factors that make the sheep model best suited for assessing fetal development. First, this model’s most notable strengths include similarities in blood gas profiles, hormone responses, organ development, and birthing profiles generally comprised of singletons or twins with weights mirroring those of human infants. Second, both ovine ewes and fetuses allow for long-term surgically implanted data collection devices, unlike other large animal models such as pigs.^7^ Finally, sheep are more feasible financially and in terms of care than other large animal models, such as non-human primates, while allowing for long-term sampling of the same animal to monitor change over time, a common drawback for smaller species models.^1^

Pregnant sheep models have paved the way for momentous discoveries such as glucocorticoid stimulated production of surfactant in fetuses at risk for respiratory distress syndrome.1 Through sheep models, researchers found that hypothermia can be used to mitigate the effects of asphyxiation during birth, established the development of intrauterine surgery, and, more recently, discovered the possibility of extracorporeal fetal development.^8^ Furthermore, researchers have successfully translated physiological information derived directly from the pregnant sheep model to standard clinical practice in human perinatal care.^1^

The pregnant sheep model provides a clear avenue to build upon these discoveries and further expand our understanding of human development. Unfortunately, there is still limited knowledge of how to properly implement the pregnant sheep model with a high degree of reliability that overcomes current barriers to these models.^9^ An issue prenatal researchers often face is ovine fetal demise, resulting in diminished study power. Another problem common to large animal models is the higher cost and significant infrastructure required to house large animal species. These issues reduce the feasibility of employing a great number of animals, which means there is minimal room for animal loss. This issue is magnified in the chronically instrumented, non-anesthetized fetal sheep model due to the inherent invasive techniques that may disturb a highly sensitive fetal environment. Current literature does not address common sources of error that cause deaths in sheep models, but research teams need this foundational information to improve the reliability of the model moving forward and to facilitate a broader adoption of this important model of human fetal physiology.

To ensure the soundness of the data produced using pregnant sheep models and the clinical guidelines derived from them, the factors that determine reproducible success must be identified and made explicit. We propose the application of systems evaluations, an approach often employed in surgery and aviation, which has strikingly diminished failure rates in these fields. We conducted a retrospective cohort analysis comprising 82 animal experiments over the course of six years to understand and identify what factors contribute to the success of a chronically instrumented non-anesthetized fetal sheep model.

## Methods

### Ethics Approval

Animal care followed the guidelines of the Canadian Council on Animal Care and the approval by the University of Montreal Council on Animal Care (protocol #10-Rech-1560).

### Cohort source

The 82 experiments were conducted at the Université de Montréal over the course of six years in studies investigating the use of fetal heart rate variability as an early biomarker for fetal inflammation and the contribution of the fetal vagus nerve. Lipopolysaccharide (LPS)-induced inflammation in fetal sheep is a well-established model of the human fetal inflammatory response to sepsis.^10,14–16^ Animals were surgically instrumented with amniotic and vascular catheters and ECG electrodes and then injected with LPS to simulate chorioamnionitis. Cytokine profiles were then compared with fetal heart rate patterns. A more detailed explanation of the methods and protocol can be found below; the full study can be found elsewhere.^2,3,10–12^

### Anesthesia and surgical procedure

We reported the detailed approach elsewhere.^2,13^

Briefly, we instrumented 82 pregnant time-dated ewes at 126 days of gestation (dGA, ~0.86 gestation) with arterial, venous and amniotic catheters, fetal precordial ECG and cervical bilateral VNS electrodes; 19 animals received cervical bilateral vagotomy (Vx) during surgery of which 8 animals received efferent and 6 animals afferent VNS electrodes and VNS treatment during the experiment.^2,13^

Ovine singleton fetuses of mixed-breed were surgically instrumented with sterile technique under general anesthesia (both ewe and fetus). In the case of twin pregnancy, the larger fetus was chosen based on palpating and estimating the intertemporal diameter. The total duration of the procedure was carried out in about 2 hours. Antibiotics were administered to the mother intravenously (Trimethoprim sulfadoxine 5 mg/kg) as well as to the fetus intravenously and into the amniotic cavity (ampicillin 250 mg). Amniotic fluid lost during surgery was replaced with warm saline. The catheters exteriorized through the maternal flank were secured to the back of the ewe in a plastic pouch. For the duration of the experiment, the ewe was returned to the metabolic cage, where she could stand, lie and eat ad libitum while we monitored the non-anesthetized fetus without sedating the mother. During postoperative recovery antibiotic administration was continued for 3 days. Arterial blood was sampled for evaluation of the maternal and fetal condition and catheters were flushed with heparinized saline to maintain patency. As reported in Burns et al.(Burns et al. 2015), care was taken to provide adequate analgesia intrasurgically as well as during the post-operative period. This is important because analgesic therapies reduce maternal stress which in turn may translate into lower fetal stress.

### Experimental protocol

Postoperatively, all animals were allowed 3 days to recover before starting the experiments. On these 3 days, at 9.00 am 3 mL arterial plasma sample was taken for blood gasses and cytokine analysis. Each experiment commenced at 9.00 am with a 1 h baseline measurement followed by the respective intervention as outlined below (Fig. 1). FHR and arterial blood pressure was monitored continuously (CED, Cambridge, U.K., and NeuroLog, Digitimer, Hertfordshire, U.K). Blood and amniotic fluid samples (3 mL) were taken for arterial blood gasses, lactate, glucose and base excess (in plasma, ABL800Flex, Radiometer) and cytokines (in plasma and amniotic fluid) at the time points 0 (baseline), +1 (*i.e*., immediately after LPS administration), +3, +6, +24, +27, +30, +48 and +54 h (*i.e*., before sacrifice at day 3). For the cytokine analysis, plasma was spun at 4°C (4 min, 4000g force, Eppendorf 5804R, Mississauga, ON), decanted and stored at −80°C for subsequent ELISAs. After the +54 hours (Day 3) sampling, the animals were sacrificed. Fetal growth was assessed by body, brain, liver and maternal weights.

**Figure 1.**
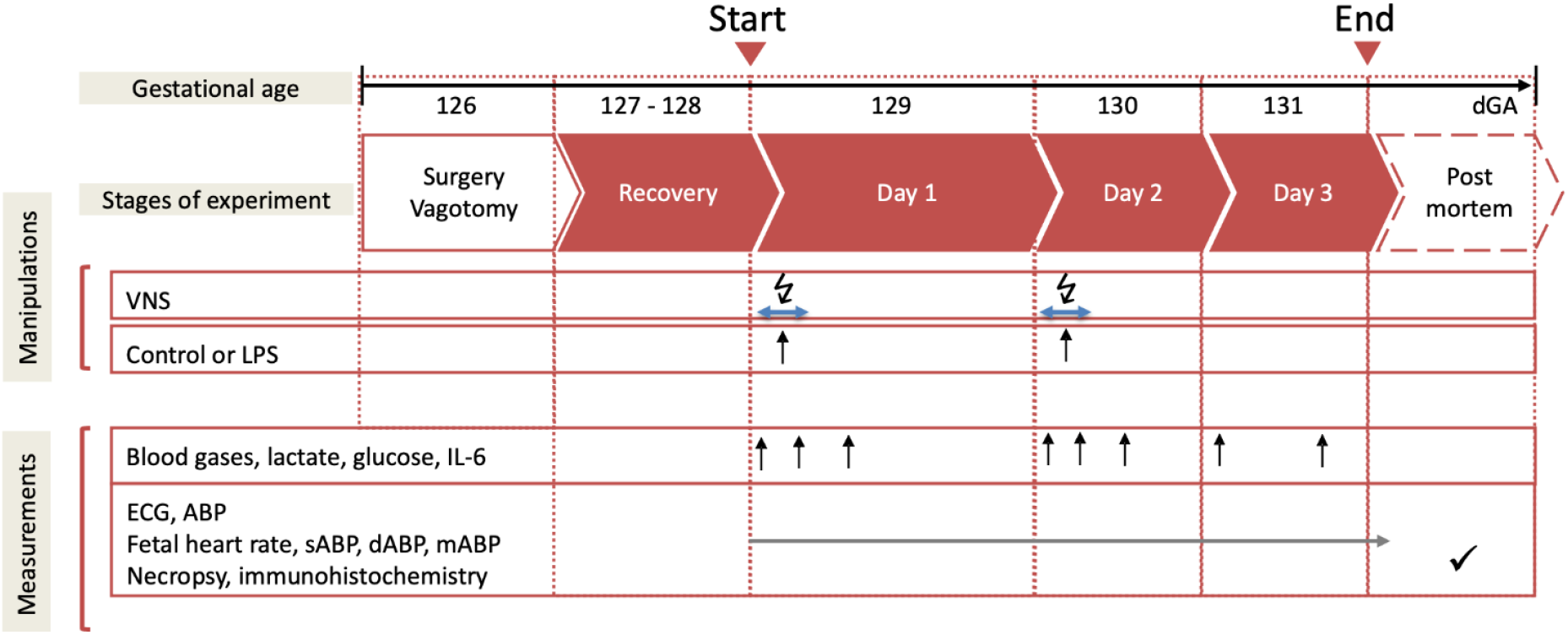
Experimental design. Bilateral cervical vagotomy (Vx) was performed during surgery in Vx+LPS group animals. At Days 1 and 2, Vx+LPS animals received LPS dose 400 or 800 ng/fetus/day. Some Vx+LPS animals also received the efferent (to the periphery) or afferent (to the brain) intermittent VNS on Days 1 and 2 (VNS groups). Reproduced from Cao et al 2022.11

The experimental groups consisted of four following categories.

#### Control and LPS groups

Fifteen fetuses were used as controls receiving NaCl 0.9%. Twenty-three fetuses received LPS (100 n=2, 200 n=1, 400 n=15 or 800 n=5 as ng/fetus/day) derived from E. coli (Sigma L5293, from E coli O111:B4, ready made solution containing 1 mg/ml of LPS) were administered intravenously to fetuses on days 1 and 2 at 10.00 am to mimic high levels of endotoxin in fetal circulation over several days as it may occur in chorioamnionitis. As we identified that IL-6 response did not depend on the LPS dose in the applied range,^10^ these animals were all considered as one LPS group for statistical comparison purposes.

#### Vx+LPS groups

Eleven animals were vagotomized (Vx) and exposed, similar to the LPS group, to LPS400 (n=6) or LPS800 (n=5).

#### VNS groups

Efferent (n=8) or afferent (n=6) VNS probes were installed, distally or proximally of the Vx site, respectively.

### Retrospective cohort analysis

To investigate factors that dictate the success of a chronically instrumented non-anesthetized fetal sheep model, ten variables (listed below) common to all 82 experiments plus the temporal order of the experiments were evaluated. We deemed a failure to be an experiment that involved the premature death of an animal or fetus for unknown reasons. In total, over six years, 28% of experiments experienced such “failure”.

- Temporal order as a proxy for the experience level of the team (encoded as the order of the outcomes)
- Time of year (season: fall, spring, summer)
- Number of fetuses
- Maternal weight
- Fetal weight (instrumented fetus)
- Gestational age at surgery
- Gestational age at euthanasia
- Surgery duration
- Length of time uterus is open during surgery, i.e., duration of fetal surgery
- Timing of first fetal blood sampling
- Vagotomy during surgery (yes/no)

We deliberately excluded instances of poor experimental outcomes with known causes due to human error. The two key factors in the early stages of the model were

1. Catheter flushing: the technique of flushing the fetal and maternal vascular catheters with saline to maintain patency. We reported the technique elsewhere.^2^
2. Pregnancy ketosis: We found it useful for monitoring ewe’s health to measure once daily dam’s ketosis status using an over-the-counter ketone bodies strip from veinous blood sample as early as during the surgical instrumentation. Our internal veterinary medicine team established a procedure to initiate corrective measures via continuous i.v. infusion when necessary during the three days of post-operative recovery.

#### Statistical approach

To evaluate the role of each variable in contributing to the success of the model, a binary logit regression analysis with a Fisher scoring optimization was fit to the data (SAS Studio, V9 engine, release 3.8). A higher predictive probability indicates a larger impact by the given variable on the outcome of the experiment. A Wald Chi-Square analysis was run on the data to control for confounds and determine significance. The raw data, the SAS code and the output were deposited on FigShare.^18^

## Results

The ten predictive variables were analyzed to investigate which factors dictate the success of a chronically instrumented fetal sheep model. A summary overview of these variables is presented in Table 1.

**Table 1.** Cohort overview.

| Seasons |  | Number of Fetuses |  |
| --- | --- | --- | --- |
| *Fall experiments* | 27 (32.9%) | 1 | 29 (35.4%) |
| *Spring experiments* | 44 (53.7%) | 2 | 48 (58.5%) |
| *Summer experiments* | 11(13.4%) | 3 | 5 (6.1%) |
| Vagotomy |  | Length of time uterus is open (min) |  |
| Yes | 32 (39%) | *Median* | 86.5 |
| No | 50 (61%) | *IQR* | 46.8 |
|  |  | Timing of 1st fetal sample (min) |  |
|  |  | *Median* | 39 |
|  |  | *IQR* | 13 |
| Maternal Weight (kg) |  | Fetal Weight (kg) |  |
| *Median* | 70 | *Median* | 3.6 |
| *IQR* | 22.9 | *IQR* | 1.7 |
| Gestational age at surgery (Days) |  | Gestational at euthanasia (days) |  |
| *Median* | 127 | *Median* | 133 |
| *IQR* | 4.5 | *IQR* | 4.7 |

Only three variables, gestational age at euthanasia (PD_dGA), number of fetuses, and temporal order were initially found to be significant (p=0.011, p<0.0001, and p<0.0001, respectively). However, upon controlling for the temporal order, the contribution of the variables PD_dGA and “number of fetuses” became non-significant (p=0.3762 and p=0.1189, respectively). Thus, the only factor that influences success of the experimental model is the level of experience the team has with implementing such a model.

## Discussion

We demonstrate that the key variable for determining the success of a chronically instrumented non-anesthetized fetal sheep model is the experience level of the team. Our findings will help improve the quality of data produced using such a model and will allow researchers to better understand the causes for model failure.

A common yet conventionally underreported issue researchers face with the pregnant sheep model is animal loss. This reduces study power and the subsequent generalizability of results. Given the time, effort, and resources often dedicated to the deployment of a pregnant sheep model, understanding the reasons behind animal loss in sheep model studies is critical.

In an effort to ascertain the root cause of animal loss, our team looked at ten variables we deemed most likely contributing to animal loss, based on our subjective experience with the model and objective understanding of the contributing causes. Interestingly, four factors initially deemed to be of the highest importance, number of fetuses, fetal weight, maternal weight, and length of time the uterus is open during surgery, had no statistical impact on the rate of animal loss. Our team predicted the number of fetuses and weights to increase the technical difficulty of the surgery, potentiating error and resulting in animal demise. We predicted that the length of time the uterus was open for instrumentation acts as both a proxy for surgical difficulty and a potential source for infection. Contrary to our assumptions, high-risk animals and surgical delays had no impact, implying the pregnant sheep model is more resilient to external intervention than initially believed. However, these four factors have a common factor that is, the abilities of the surgical and post-operative teams to handle such challenges grow with experience.

Of the ten variables investigated, only temporal order showed a statistically significant relationship with the success rate of the model. Temporal order was used as a proxy for the experience level of the team. Experience level plays a pivotal role because adherence to standard operating procedures (SOP) improves with time ensuring best practices, and ultimately best outcomes. Communication and problem-solving capabilities also improve as team members expand their working knowledge of the model. The data indicates that experienced surgeons will more closely follow protocols and, when problems arise, better navigate issues associated with factors such as low weights or multiparity. These findings carry into post-operative care as well due to a more robust awareness of infection prevention, medication administration, and wound care. As such, the magnitude of risks associated with a pregnant sheep model is not intrinsic to the model itself but relative to the research team, including the experience level of the surgeon to the level of the technicians conducting post-operative care.

Our findings echo a long-standing notion in high-risk environments – more experienced teams perform better. A study investigating causes for poor surgical outcomes in humans found that the majority of factors were related to attributes developed through practice. The researchers found poor communication, information overload, poor management of stress, deviations from standard practice, interruptions, distractability, hierarchical structures, and inadequate skill to be at the root of many poor surgical outcomes in humans.^19^ We feel many of these same factors are what drives team experience to dictate the success in the chronically instrumented non-anesthetized fetal sheep model of human development. The issue of “deviation from standard practices” as addressed earlier, is one factor our team has attempted to combat by publishing a SOP in 2015.^2^ However, even with a clear SOP, improper execution of the procedures is a common source of error leading to surgical complications, infections, and medication errors. We believe this stems from poor intra-team communication, improper training, and a lack of clear roles/responsibilities. Training is an obvious component of experience, but it is also easily overlooked. Assumptions of prior understanding or notions about hierarchical role importance can lead to poor training. These misunderstandings are potential causes of error as the experience level of all team members dictates model success – not only those team members at the “top.” For example, unclear expectations from team leaders can cause dosing errors for medications like antibiotics or heparin, or by preventing foresight of issues that may arise from an overburdened, absent, or confused team member. To overcome such challenges we suggest the implementation of closed-loop communication styles. This is a practice where orders are repeated, out loud, back to the person giving them. Fields such as surgery have found great success in preventing errors that arise from miscommunication or inexperience by implementing closed-loop communication. These practices in the setting of the fetal sheep model would benefit not only new teams but even the most experienced groups.

In addition to training and improving intra-team communication, we feel improved transparency of failures can serve as a potential avenue for increased success. Inter-team collaboration prevents repeating challenges other research teams have already overcome, thereby increasing experience levels without requiring each team to endure the same tribulations. Currently, it is rare to openly publish data highlighting the time and resources spent fine-tuning models. However, our data indicate that withholding such experience is detrimental to science and study development. Experience level is more than simple adherence to an SOP, it is the ability to predict errors, solve problems, and ensure everyone is working together. Error prediction and problem-solving are two skills typically gained through trial and error and learning from previous mistakes. We propose streamlining this learning process through increased inter-team collaboration. Working as a field to build off and learn from one another’s studies will improve everyone’s abilities to derive reliable data from a pregnant sheep model.

There are limitations to mention. First, by setting up the ten causally hypothesized contributing factors that determine the outcome of the experiment, we made exclusive assumptions. There may be other predictors, not accounted for in this model, beyond the temporal order, that could determine the outcome. Second, we did not explicitly take into account the incremental improvements in anesthesia. We do not estimate these improvements to be a strong contributor to outcomes for the entire cohort, because these occurred early in the course of the experiments. Nonetheless, we wish to emphasize the importance of a strong anesthesia team in this large animal experimental model that is paying attention to the well-being of the ewe with regard to tocolytics and pain relief. Because of the unique characteristics of the model, any stress-relieving interventions on the mother are likely to positively affect the fetal well-being as well.

To continue expanding the realm of possibility for prenatal medicine, research teams must understand what dictates the success of a pregnant sheep model. Errors in the methods of data collection will prevent even the most promising studies from achieving their full potential. Our data indicates the complexity of the study does not necessarily dictate outcomes, but, rather, the adherence to SOPs, knowledge base, and hands-on experience level of the research team conducting it impacts sheep model outcomes. These variables can be improved through robust training, collaboration, and communication. Our findings highlight the need for a shift in culture where both success and failures are openly discussed in the scientific community. Allowing research teams to learn from others’ experiences functions as the simplest and most effective path to increased success of the chronically instrumented non-anesthetized fetal sheep model of human development.

## Acknowledgements

The authors gratefully acknowledge technical support by Esther Simard, Marco Bosa, Pierre-Yves Mulon and L. Daniel Durosier during the experiments. The authors thank the amazing Clinical Sciences team of the Faculté médecine vétérinaire at the Université de Montréal for the skilful assistance throughout the experiments. The authors also gratefully acknowledge the statistical support by Miguel Chagnon.

